# Structural Organization of the Nvj3-Mdm1 Complex Reveals a Conserved Lipid-Compatible Contact Site Module

**DOI:** 10.64898/2026.06.29.735323

**Authors:** Marwa Aboumourad, Hanaa Hariri

## Abstract

Membrane contact sites are organized by protein assemblies that physically couple organelles and coordinate lipid metabolism, yet the structural principles that enable lipid exchange across these junctions remain poorly defined. At the nuclear-vacuolar junction (NVJ) in budding yeast, the tethering protein Mdm1 and its binding partner Nvj3 form a complex that regulates lipid metabolic pathways, but the structural features underlying their interaction have not been resolved. Here, we use AlphaFold-based complex prediction and comparative structural analysis to define the organization of Nvj3-Mdm1 complex assembly. We identify a high-confidence heterodimer in which conserved PXA and PXC domains generate an extended tunnel spanning both proteins. Tunnel analysis predicts a core hydrophobic conduit traversing the Nvj3-Mdm1 interface, consistent with a lipid-compatible architecture. Evolutionary conservation is enriched at the Nvj3-Mdm1 interface. The predicted conduit shares geometric and physicochemical properties with bridge-like lipid transfer proteins, including Atg2, Fmp27, and Hob2, suggesting that heteromeric tether assemblies may contribute directly to inter-organelle lipid transfer. Notably, this conduit is predicted to arise from a heteromeric α-helical assembly rather than the β-sheet-rich architecture characteristic of canonical bridge-like lipid transfer proteins. Comparative phylogenetic analyses showed that Nvj3 and Mdm1 share broadly congruent evolutionary patterns across Saccharomycetes, consistent with their conserved functional association. Together, these findings define Nvj3 as a structural partner of Mdm1 and support a conduit-based model of lipid transfer at the NVJ.

## Introduction

Membrane contact sites are regions where organelles come into proximity to coordinate lipid exchange, metabolite transfer, and cellular homeostasis (Bohnert, 2020; Hariri, 2021; Prinz et al., 2020; Scorrano et al., 2019a). The nuclear-vacuolar junction (NVJ) in *Saccharomyces cerevisiae* represents a prominent endoplasmic reticulum (ER)-vacuole contact site that plays central roles in lipid metabolism, including the regulation of lipid droplets (LDs) and stress-responsive membrane remodeling (Hariri et al., 2018, 2019; Henne et al., 2015; Kvam & Goldfarb, 2006; Obaseki et al., 2024; Pan et al., 2000). Despite their functional significance, the structural mechanisms that organize NVJs and enable lipid flux across this junction remain incompletely understood. In particular, whether NVJ tether proteins merely position metabolic enzymes or directly contribute to lipid transfer is unknown.

Mdm1 and Nvj3 are core components of the NVJ, with Mdm1 functioning as an ER-anchored tether and Nvj3 implicated in coordinating lipid metabolic pathways (Adebayo et al., 2026; Hariri et al., 2019; Henne et al., 2015). Previous studies identified Nvj3 as an interacting partner of Mdm1, and recruitment of Nvj3 to the NVJ has been shown to depend on Mdm1 (Henne et al., 2015). While Nvj3 is predominantly cytosolic under basal conditions, it localizes to the NVJ under metabolic stress conditions, suggesting that its recruitment to this contact site is dynamically regulated (Adebayo, Obaseki, Miller, et al., 2026).

Mdm1 belongs to the conserved SNX-RGS family of sorting nexins and contains an N-terminal ER-anchoring region, a central Phox-homology (PX) domain that mediates membrane association, and flanking PX-associated (PXA and PXC) domains (Chandra et al., 2019; Hariri & Henne, 2022). Nvj3 shares a related domain organization, including predicted PXA and PXC domains, although its functional architecture remains less well characterized. These features suggest that both proteins may possess lipid-interacting surfaces and could assemble into a structurally organized interface at the NVJ. Despite this evidence, the molecular basis for Nvj3-Mdm1 interaction and how this complex contributes to NVJ organization remain poorly understood.

Both proteins contain predicted hydrophobic domains and internal cavities, raising the possibility that their association could generate a lipid-compatible interface (Castro et al., 2022; Paul et al., 2022a). Structural predictions suggest that Mdm1 contains a hydrophobic cavity reminiscent of lipid transfer proteins, supporting a potential role in inter-organelle movement during metabolic stress. Nvj3 exhibits similar architectural features, yet whether these proteins assemble into a complex or instead function completely independently has not been fully explored. Consequently, the molecular basis by which Mdm1 and Nvj3 cooperate to shape NVJ architecture and lipid organization remains unresolved.

Recent advances in computational structural biology allow for high-confidence prediction of protein complexes and the identification of internal channels that may support lipid transfer (Neuman et al., 2022; Levine, 2022). In Atg2, structural and biochemical work showed that the conserved N-terminal region contains a hydrophobic cavity that accommodates phospholipid acyl chains, bridges membranes, and supports direct phospholipid transfer in vitro (Osawa et al., 2019). Related Vps13-like proteins also function as bridge-like lipid transport systems; for example, Csf1 supplies phosphatidylethanolamine to the ER for GPI-anchor synthesis, supporting a physiological role for tube-like lipid transport proteins in glycerophospholipid delivery (Toulmay et al., 2022). More recent structural and simulation-based studies further suggest that lipids can move through these hydrophobic conduits in an organized manner, reinforcing the model of bridge-like proteins as membrane-spanning lipid-transfer systems (Wang et al., 2025). Notably, a similar bridge-like mechanism has also been described in bacteria, where studies on YhdP show lipid transfer through an extended hydrophobic groove spanning the Gram-negative cell envelope, from the inner membrane across the periplasm to the outer membrane (Cooper et al., 2025).

Here, we use AlphaFold to model the Nvj3-Mdm1 heterodimer and uncover a continuous hydrophobic tunnel spanning their interface. We further characterize the geometry, physicochemical properties, and evolutionary conservation of this feature and compare it with known bridge-like lipid-transfer proteins (BLTPs). Additionally, co-phylogenetic analysis is consistent with shared phylogenetic history between Nvj3 and Mdm1 across Saccharomycetes. Our findings support a model in which the Nvj3-Mdm1 assembly forms a structurally integrated platform that may facilitate lipid movement, providing a foundation for future mechanistic investigation of tether-mediated lipid transport at the NVJ.

## Results

### AlphaFold predicts a heterodimeric Nvj3-Mdm1 complex assembly

The Nvj3-Mdm1 complex has been implicated in maintaining NVJ integrity and regulating lipid-associated processes; however, the molecular organization of this complex and how it contributes to NVJ function remains unclear. In particular, whether the structural arrangement of Nvj3 and Mdm1 could support lipid transfer across the junction has not been explored. Because AlphaFold-based complex prediction does not by itself establish biological interaction, we used AlphaFold3 to evaluate the structural plausibility, organization, and interface features of the previously implicated Nvj3-Mdm1 complex.

Prior work and domain predictions suggest that both proteins harbor internal hydrophobic cavities (Paul et al., 2022b), suggesting that their association may generate a structurally organized hydrophobic interface.

Recent breakthroughs in computational protein structure prediction, specifically the development of AlphaFold, have made it possible to model proteins and their complexes at near-atomic resolution (Abramson et al., 2024). AlphaFold3 predicted an elongated heterodimeric assembly in which Nvj3 and Mdm1 form an extended complex composed predominantly of α-helical domains connected by flexible linkers (Figure 1A), where the structure is colored according to predicted local distance difference test (pLDDT) scores. The majority of structured regions are predicted with high confidence (pLDDT 70-90), while terminal regions display lower confidence consistent with flexible or intrinsically disordered segments. The major predicted domains were mapped onto the structural model to clarify the organization of Mdm1 and Nvj3 within the complex, including the Mdm1 transmembrane region, PXA, PX, and PXC domains, and the Nvj3 PXA and PXC domains (Supplementary Figure 1).

**Figure 1:**
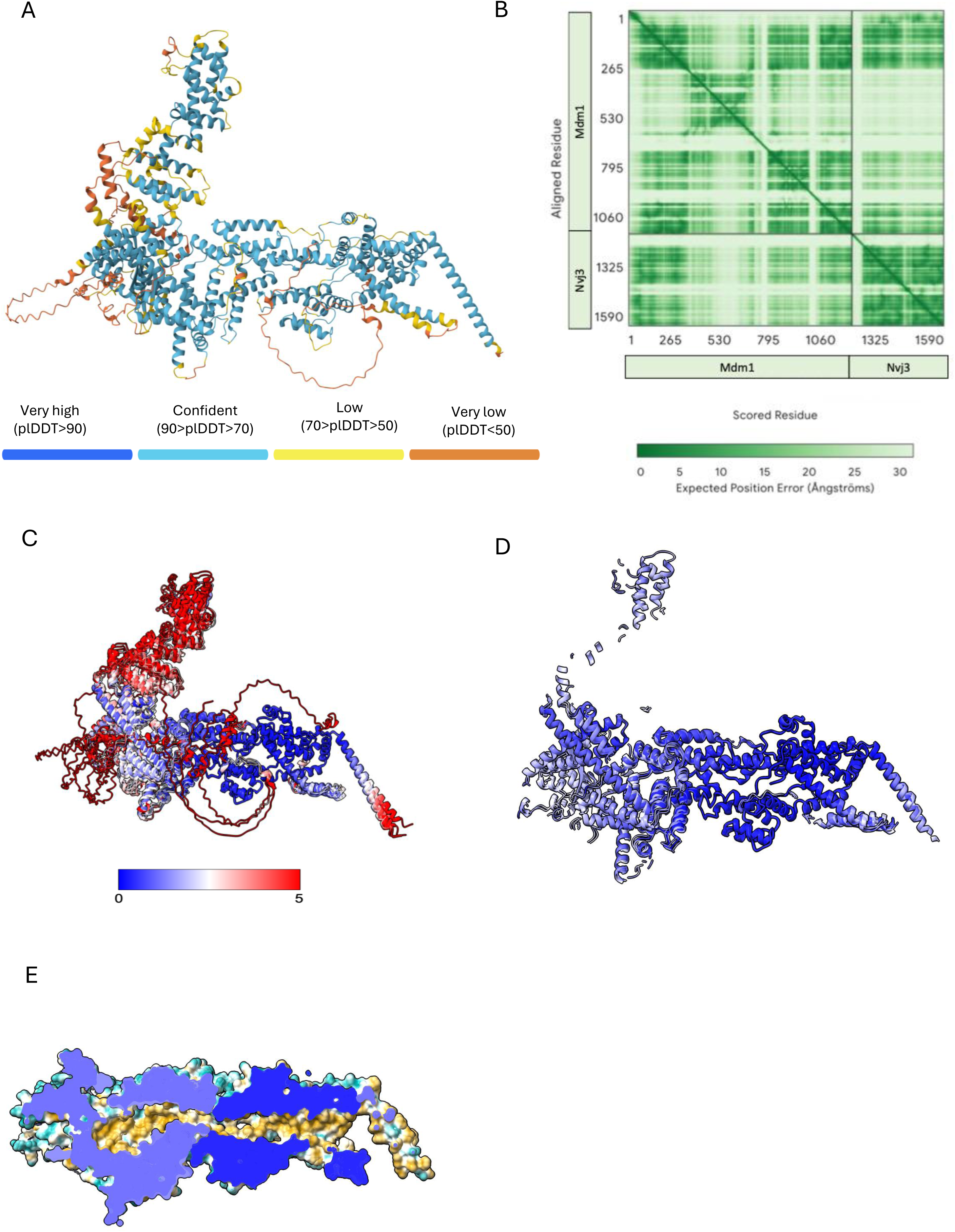
Structural Superposition of AlphaFold3 Models of the Nvj3-Mdm1 and analysis of stability and interface. A) Alphafold predicted structure of the Nvj3-Mdm1 complex shown as a cartoon and colored by per-residue confidence (pLDDT), where blue indicates high confidence and yellow/orange indicate lower confidence. B) Predicted Aligned Error (PAE) plot showing the expected positional error between residue pairs across the complex, where darker colors indicate higher confidence in the relative positioning. C) Superposition of AlphaFold models 0-4 of the Nvj3-Mdm1 complex rendered in ribbon representation and colored by per-residue RMSD (0-5 Å). Deep blue regions indicate structurally conserved rehions, whereas bright red indicates conformational displacement > 5 Å. D) Visualization of the stable structural core across all models, with flexible residues (RMSD ≥ 2 Å) removed. E) Side-view cross-section of the stable core interface from Panel D, colored by surface hydrophobicity.

To further evaluate model reliability, we examined the predicted aligned error (PAE) matrix (Figure 1B). The PAE map reveals low intra-chain positional error within both Mdm1 and Nvj3, visible as the darker green blocks along the main diagonal corresponding to each protein. In this context, intra-chain positional error reflects the predicted uncertainty in the relative positioning of the residues within the same polypeptide chain; low values indicate that different regions of the protein are confidently positioned relative to one another, supporting a well-defined domain architecture. The off-diagonal regions corresponding to interactions between Mdm1 and Nvj3 indicate a degree of flexibility in their relative positioning. This suggests that while each protein adopts a well-defined structure, the overall arrangement of the complex may allow for conformational variability.

To assess structural robustness, the five ranked output models by AlphaFold3 were compared by structural superposition and colored by root-mean-standard deviation (RMSD), which measures the average distance between corresponding atoms in aligned structures and provides a quantitative indicator of structural similarity (Figure 1C, Table S1). The RMSD color scale was capped at 5 Å, with higher values displayed in red to highlight regions of increased structural variability, while blue denotes structurally conserved regions. Global RMSD values reached up to 22.129 Å, reflecting substantial conformational variability in peripheral regions (Table S1). In contrast, after excluding highly variable segments with RMSD ≥ 5 Å, the central conduit-forming core showed low RMSD values ranging from 0.438 to 1.091 Å (Figure 1D; Table S1). This supports strong structural agreement across the AF3 output models. These results indicate that conformational differences arise primarily from flexible terminal domains and linker regions, while the central Nvj3-Mdm1 binding interface remains structurally stable. The high-variability segments likely correspond to intrinsically disordered regions, which may provide the flexibility needed to accommodate the dynamic spatial organization of the NVJ.

To further quantify inter-chain confidence at the predicted Nvj3-Mdm1 interface, we extracted numerical PAE values for selected interfacial residue pairs using the AlphaFold3 PAE output and the ChimeraX alphafold contacts command. The analyzed inter-chain contacts showed PAE values ranging from 3.60 to 7.20 Å, with a mean interfacial PAE of 5.06 Å supporting high-confidence relative positioning of the selected Nvj3-Mdm1 interface contacts (Supplementary Table 3).

Further inspection of the interaction interface using a side-view cross-section revealed a predominantly hydrophobic interior (Figure 1E), with detailed characterization provided in Figure 2. To evaluate whether the predicted interface corresponds to a biophysically favorable interaction, we analyzed binding energetics using the PRODIGY server (PROtein binDIng enerGY) (Xue et al., 2016). PRODIGY estimates binding free energy (ΔG) and dissociation constant (Kd) based on structural features, where more negative ΔG values correspond to stronger and more favorable interactions, and lower Kd means tighter binding. This analysis predicted a highly stable complex (ΔG = −15.1 kcal·mol⁻¹, Kd ≈ 8.9 × 10⁻¹² M at 25 °C). The interface comprises 160 intermolecular contacts, dominated by 55 apolar-apolar and 38 charged-apolar interactions, indicating that hydrophobic interactions dominate the binding interface (Table 1).

**Figure 2:**
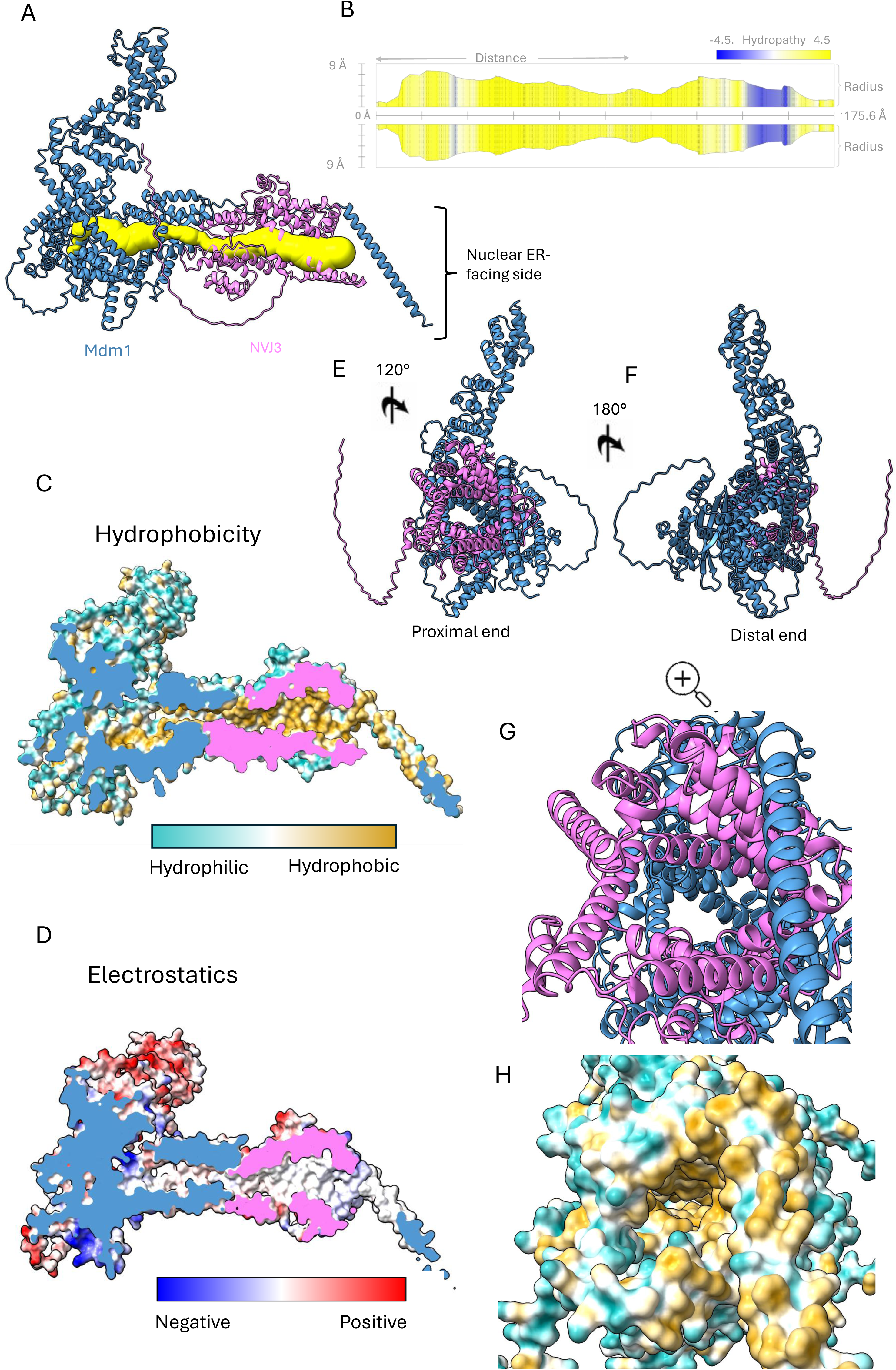
Nvj3-Mdm1 heterodimeric interaction and tunnel characterization. A) Cartoon representation of the predicted Nvj3-Mdm1 complex, with Mdm1 colored in blue and Nvj3 colored in pink. The internal tunnel is highlighted in yellow, identified using MOLE online. The bracket indicates the nuclear ER-facing side, where the Mdm1 transmembrane helix is located. B) Tunnel profile along its length, showing a 175.6 Å continuous pathway and corresponding hydropathy distribution, consistent with a predominantly lipophilic environment. C) Surface representation colored by hydrophobicity, shown as a longitudinal cross-section to reveal the tunnel interior, which exhibits a continuous lipophilic character. D) Surface representation colored by electrostatic potential, showing a largely neutral interior within the tunnel lining. E) Alternative view of the Nvj3-Mdm1 complex rotated 120° relative to the orientation in panel A, highlighting the overall spatial organization and exposing the tunnel entrance at the proximal, nuclear ER-facing end of the complex. F) Alternative view of the Nvj3-Mdm1 complex rotated 180° relative to the orientation in panel E, showing the distal end of the complex. This end lies opposite the proximal nuclear ER-facing side and corresponds to the vacuolar-binding side. G) Zoomed-in view of the proximal tunnel entrance shown in panel E. H) Zoomed-in view of the proximal tunnel entrance colored according to hydrophobicity, showing a lipophilic interior lining consistent with accommodation of lipid acyl chains.

**Table 1.** PRODIGY-based interface analysis of the Nvj3-Mdm1 complex.

Together, these results support a structurally plausible model of the Nvj3-Mdm1 complex with a hydrophobic interaction interface at the NVJ.

### A continuous hydrophobic tunnel connects the Nvj3-Mdm1 interface

To determine whether the structural organization of the Nvj3-Mdm1 complex could theoretically support lipid movement at the nucleus-vacuole junction, we examined the internal architecture of the predicted heterodimeric interface. Using the MOLEonline server, which identifies channels by mapping the accessible void space within protein structures and tracing continuous pathways through it (Pravda et al., 2018), we identified a continuous tunnel spanning the interface between the PXA and PXC domains of Mdm1 (blue) and complementary regions of Nvj3 (pink) (Figure 2A, Supplementary Figure S1). The tunnel extends approximately 175.6 Å across the dimer interface and exhibits openings at both ends. One opening is positioned near the Mdm1 ER-associated transmembrane region, where Nvj3 is also predicted to associate, while the opposing opening lies on the opposite face of the complex near the vacuole-binding PX domain of Mdm1, consistent with a continuous conduit traversing the interface of the Nvj3-Mdm1 complex and aligned with the membrane contact site (Figure 2A, yellow; Movie 1).

Hydropathy analysis revealed a uniformly lipophilic environment along the predicted tunnel axis, enriched with nonpolar residues including leucine, isoleucine, valine, and phenylalanine, with relatively few polar or charged residues (Figure 2B; Supplementary Table S2). A longitudinal cross-section along the Z-axis further shows a continuous hydrophobic core extending through the interface (Figure 2C; Movie 2), while electrostatic mapping indicates that this region is largely neutral, lacking pronounced charge relative to the surrounding protein surface (Figure 2D).

Rotated views of the complex further clarify the orientation of the tunnel openings. A 120° rotation of the complex along the Y-axis, relative to the orientation shown in Figure 2A, exposes the proximal, nuclear ER-facing tunnel entrance (Figure 2E). A 180° rotation shows the opposite distal, vacuole-facing side, where the tunnel opening appears more constricted (Figure 2F). A zoomed-in view of the proximal entrance shows that the channel opening is surrounded by helices from both proteins (Figure 2G), and hydrophobicity coloring of the same view highlights a nonpolar interior lining the tunnel, emphasizing its accessibility and compatibility with lipid acyl chains (Figure 2H).

To further visualize whether lipid-like molecules could theoretically occupy this cavity, we performed an additional AlphaFold3 prediction of the Nvj3-Mdm1 complex in the presence of 50 copies of oleic acid (OLA), used as a representative lipid-like ligand. OLA molecules were predicted to localize along the hydrophobic cavity/interface of the complex, supporting the lipid-compatible character of the conduit (Supplementary Figure 2). Because AlphaFold-based ligand placement has not been experimentally validated for this system, this analysis is presented as a qualitative visualization of theoretical lipid accommodation rather than evidence of lipid-transfer activity.

Together, these analyses identify a continuous hydrophobic conduit spanning the Nvj3-Mdm1 interface, consistent with a lipid-compatible architecture.

### The Nvj3-Mdm1 tunnel exhibits features of lipid transfer proteins

To evaluate whether the structural properties of the Nvj3-Mdm1 tunnel are consistent with known features of non-vesicular lipid transporters, we compared the geometry and chemical characteristics of its predicted channel with those of established lipid transfer proteins (LTPs). Several membrane contact site proteins, including Atg2, Fmp27, and Hob2, exhibit elongated hydrophobic channels that bridge opposing membranes to mediate non-vesicular lipid transport (Osawa et al., 2019; Maeda et al., 2020; Qian et al., 2021;Toulmay et al., 2022). In the structural comparison, the Nvj3-Mdm1 complex displays an extended internal tunnel that is visually similar to the continuous channels observed in Atg2, Fmp27, and Hob2 (Figure 3A). However, unlike these bridge-like LTPs, which are dominated by extended β-sheet architectures, Nvj3-Mdm1 forms its predicted tunnel within a largely α-helical heteromeric assembly. Quantitative analysis supported this visual similarity, the predicted Nvj3-Mdm1 channel (175.6 Å) was shorter than the channels of Atg2 (213 Å), Fmp27 (238.8 Å), and Hob2 (302.2 Å), but it retained a comparable average diameter (12.4 Å), similar to Atg2 (12.9 Å), Fmp27 (15.64 Å), and Hob2 (14.6 Å) (Table 2).The conduit also has a strongly nonpolar interior, with a hydropathy value of 2.41, similar to those observed for Fmp27 (2.62) and Hob2 (2.33), and moderately lower than that of Atg2 (3.07) (Table 2). Together, these comparisons indicate that, despite its distinct α-helical architecture and shorter overall length, Nvj3-Mdm1 preserves key geometric and physicochemical features associated with lipid-transfer pathways.

**Figure 3:**
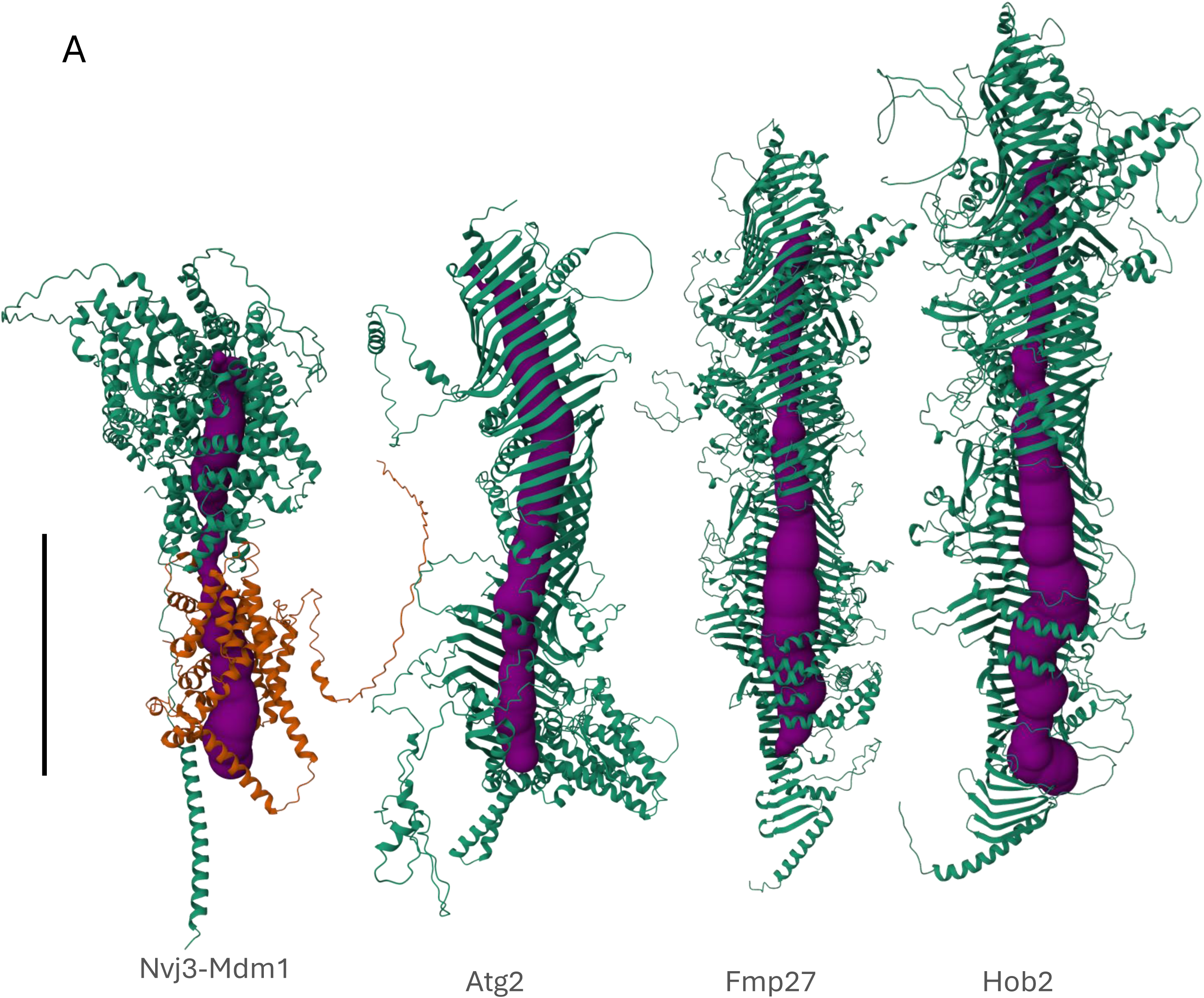
Similarity of tunnel architecture among Nvj3-Mdm1 and bridge-like lipid transport proteins. A) Structural models of the *S. cerevisiae* Nvj3-Mdm1 complex (Mdm1: green, Nvj3: orange), Atg2, Fmp27, and Hob2 are shown in cartoon representation. Predicted internal tunnels were identified using MOLEonline and are shown in purple. Scale bar = 100 Å.

**Table 2.** Quantitative comparison of tunnel length, average width, and hydropathy for the Nvj3-Mdm1 complex and representative lipid transport proteins (Atg2, Fmp27, and Hob2).

### Evolutionarily conserved residues define the Nvj3-Mdm1 interface

Residues that mediate stable and specific protein-protein interactions are often subject to strong evolutionarily constraint and tend to cluster within functional interfaces, highlighting positions critical for binding and structural stability (Guharoy & Chakrabarti, 2010). Therefore, the structural similarity between the Nvj3-Mdm1 tunnel and established lipid transfer proteins raised the question of whether this interface is evolutionarily conserved. To test this, we mapped residue-level conservation using the ConSurf algorithm as described in the Methods (Ashkenazy et al., 2016). This approach employs a standardized computational pipeline to calculate the evolutionary rate of each residue and assigns conservation grades ranging from 1 (most variable, cyan) to 9 (most highly conserved, magenta) (Figure 4).

**Figure 4:**
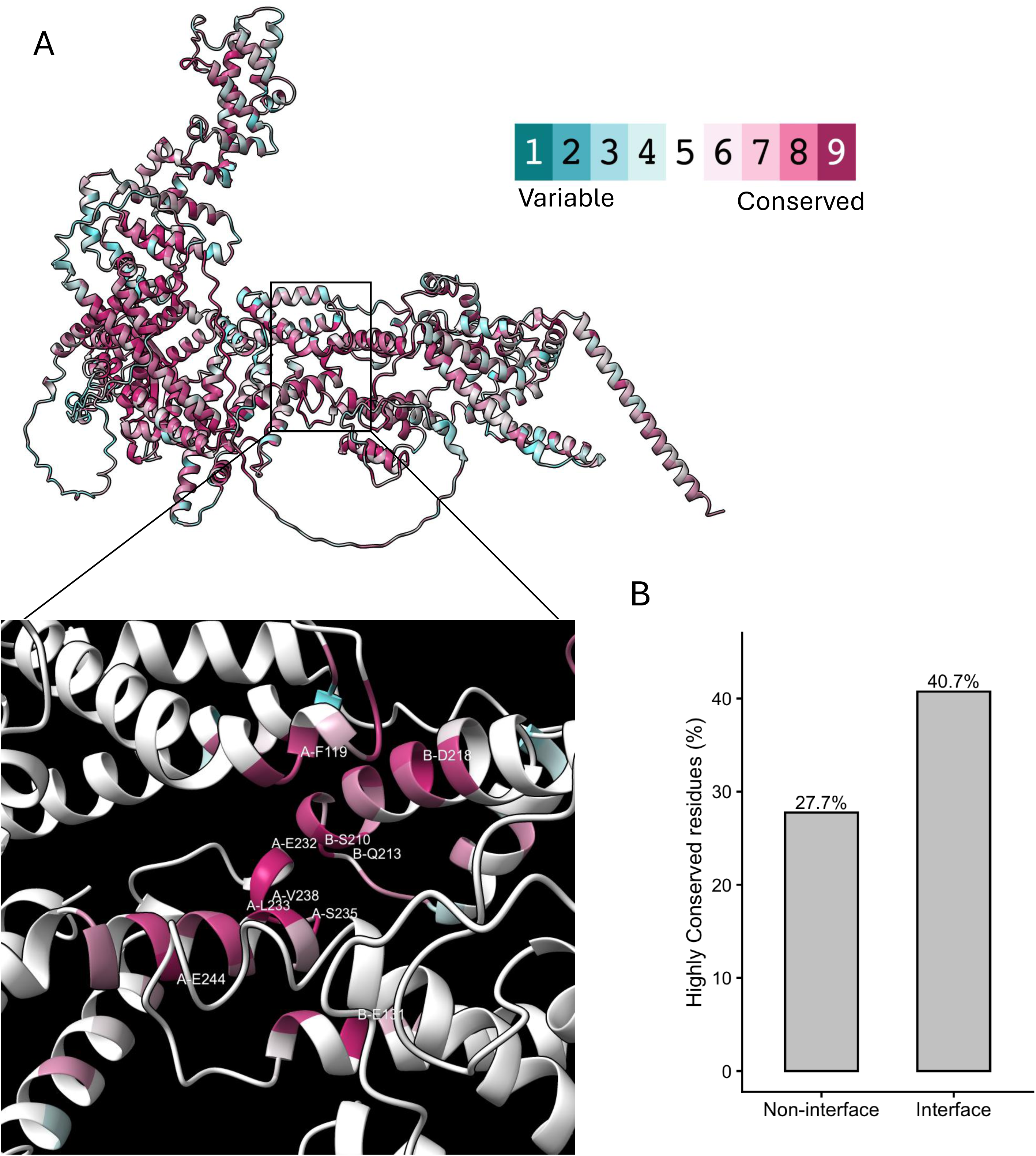
Evolutionary conservation mapping of the Nvj3-Mdm1 complex. A) Structural model of the Nvj3-Mdm1 complex colored by residue conservation scores calculated using ConSurf. The boxed region indicates the interaction interface and the inset shows a zoomed-in view with labeled residues. For clarity, only selected interface residues with the highest ConSurf conservation score of 9 are labeled. Labels indicate chain identity, with “A” corresponding to Mdm1 and “B” corresponding to Nvj3, followed by the residue identity using the one-letter amino acid code and residue position. Background ribbon elements are shown in white for visual clarity and do not indicate conservation scores. B) Quantification of conservation at the interface compared to non-interface regions. Interface residues show a higher proportion of highly conserved (score = 8 or 9) positions (40.7%) relative to non-interface residues (27.7%). This enrichment is statistically significant (Fisher’s exact test, *p* = 0.0448 indicating a trend toward increased conservation at the interaction interface).

Mapping conservation scores onto the Nvj3-Mdm1 model revealed that conserved residues are present in multiple regions throughout the complex (Figure 4A). To determine whether conservation is concentrated at the interaction interface, we examined this region in greater detail. A magnified view of the interface (Figure 4A, Inset) revealed a patch of highly conserved residues along structural elements that mediate direct inter-chain contacts, indicating a localized concentration of highly conserved residues at the interaction surface. For clarity, only interface residues are colored according to conservation, while the surrounding structure is shown in white (Figure 4 Inset).

To quantify this pattern, residues in both proteins (Mdm1 and Nvj3) were classified as interface or non-interface as described in Methods, and the proportion of highly conserved residues (ConSurf grades 8-9) was compared between these groups. Interface residues exhibited a higher percentage of conservation than non-interface residues, and this enrichment was statistically significant (Fisher’s exact test, p = 0.04) (Figure 4B), consistent with evolutionary constraint acting on residues important for interaction stability and specificity.

### Comparative phylogenetic analysis supports broad phylogenetic congruence between Nvj3 and Mdm1

Because the predicted Nvj3-Mdm1 interface showed structural organization and enrichment of conserved residues, we next asked whether this association is also reflected at the evolutionary level. Proteins that function together over long evolutionary timescales often show broadly similar phylogenetic patterns, either because they are maintained within the same species lineages or because their functional association imposes shared evolutionary constraints. Therefore, we compared Mdm1 and Nvj3 phylogenies across 120 Saccharomycetes species to determine whether their evolutionary histories are congruent. This analysis was not intended to prove direct protein-specific coevolution, but rather to provide evolutionary context for the conserved structural and functional association between Nvj3 and Mdm1.

Phylogenetic trees were constructed seperately for Mdm1 and Nvj3 using orthologous sequences identified from OrthoDB (Kuznetsov et al., 2022). We then performed cophylogenetic analysis to assess the similarity between the two trees. Visual comparison of the phylogenies revealed similar topologies, with many corresponding taxa connected by parallel or minimally crossing links. Localized crossings indicate minor differences in the relative placement of some species between the two phylogenies, which may reflect lineage-specific divergence or differences in evolutionary rate between Mdm1 and Nvj3. Overall, the parallel structure supports broad phylogenetic congruence between Mdm1 and Nvj3 across Saccharomycetes (Figure 5).

**Figure 5:**
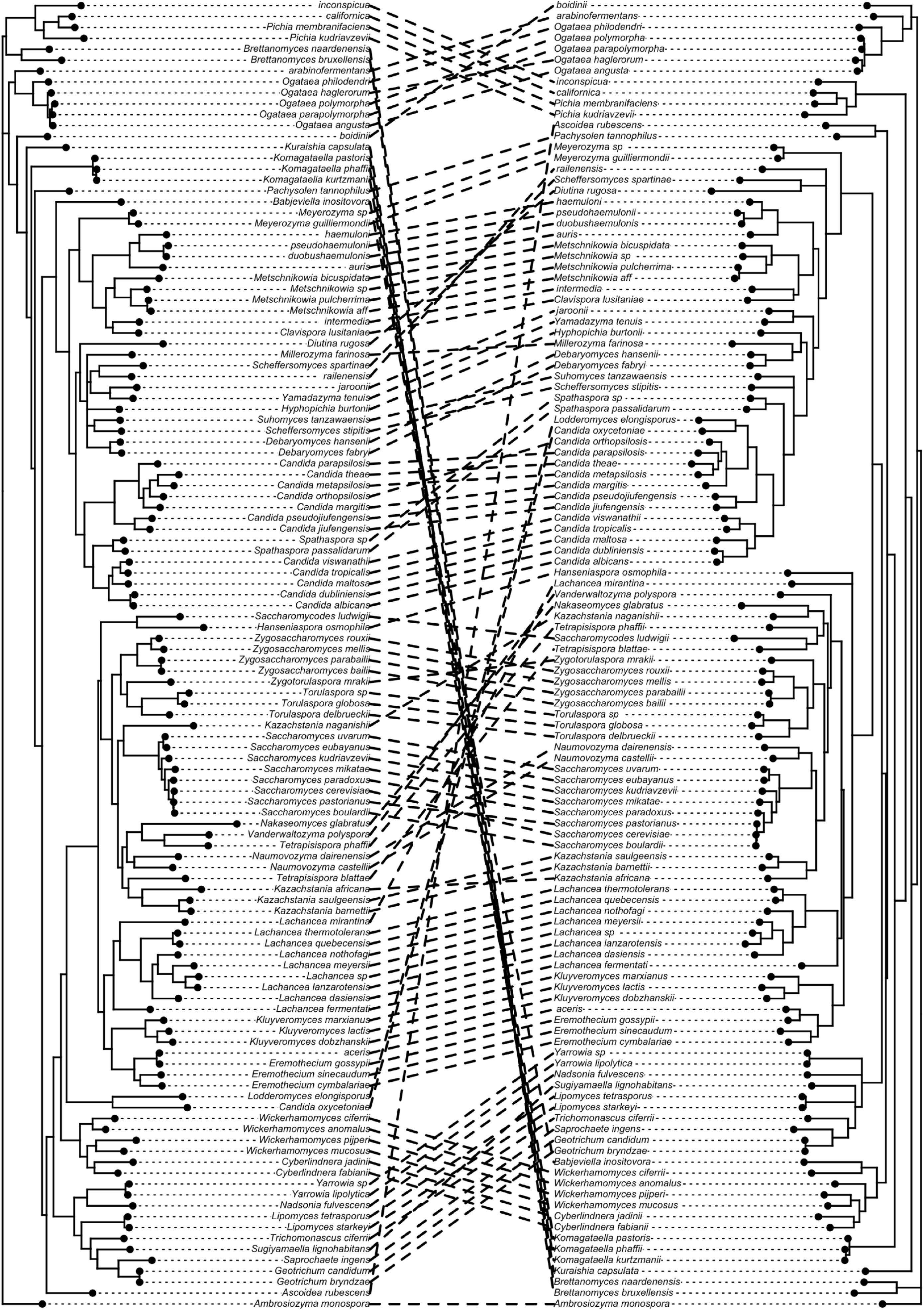
Cophylogenetic analysis across 120 Saccharomycetes species supports coordinated evolution of Mdm1 and Nvj3. Phylogenetic trees of Mdm1 (left) and Nvj3 (right) were reconstructed independently from shared Saccharomycetes species using Bayesian inference and visualized as a cophylogeny plot. Corresponding taxa are connected by dashed lines. PACo supported significant congruence between the two phylogenies (*p* < 0.001).

To quantitatively assess this pattern, we applied the Procrustes Approach to Cophylogeny (PACo) (Balbuena et al., 2013), which measures the degree of similarity between two phylogenies by comparing their pairwise evolutionary distance matrices in a shared multivariate space and assessing the goodness-of-fit between them. Statistical significance is determined by permutation testing, which evaluates whether the observed congruence exceeds that expected by chance. This analysis revealed a highly significant global congruence between the Mdm1 and Nvj3 trees (p < 0.001).

Together, these results show that Mdm1 and Nvj3 phylogenies retain broadly similar evolutionary patterns across Saccharomycetes. This congruence complements the structural conservation observed at the predicted interface and supports the inclusion of Nvj3-Mdm1 within a conserved evolutionary framework.

### The Nvj3-Mdm1 tunnel architecture is preserved in representative species

Given this significant evolutionary association, we next asked whether the hydrophobic tunnel identified in the *S. cerevisiae* represents a broadly conserved architectural feature. To address this, we selected representative Nvj3-Mdm1 ortholog pairs from the same Saccharomycetes dataset used for cophylogenetic analysis, spanning major evolutionary branches, including *Kluyveromyces lactis* and *Yarrowia lipolytica*, using the same AlphaFold-based approach followed by MOLEonline. In both species, we observed elongated internal cavities positioned near membrane-proximal regions (Figure 6A, B). Despite variation in sequence length and peripheral regions, these cavities formed continuous channels aligned along a similar axis and exhibited predominantly hydrophobic interiors. The predicted tunnel lengths were 151.2 Å in *K. lactis* and 147.4 Å in *Y. lipolytica*, indicating that extended hydrophobic cavities are retained across divergent Saccharomycetes representatives.

**Figure 6:**
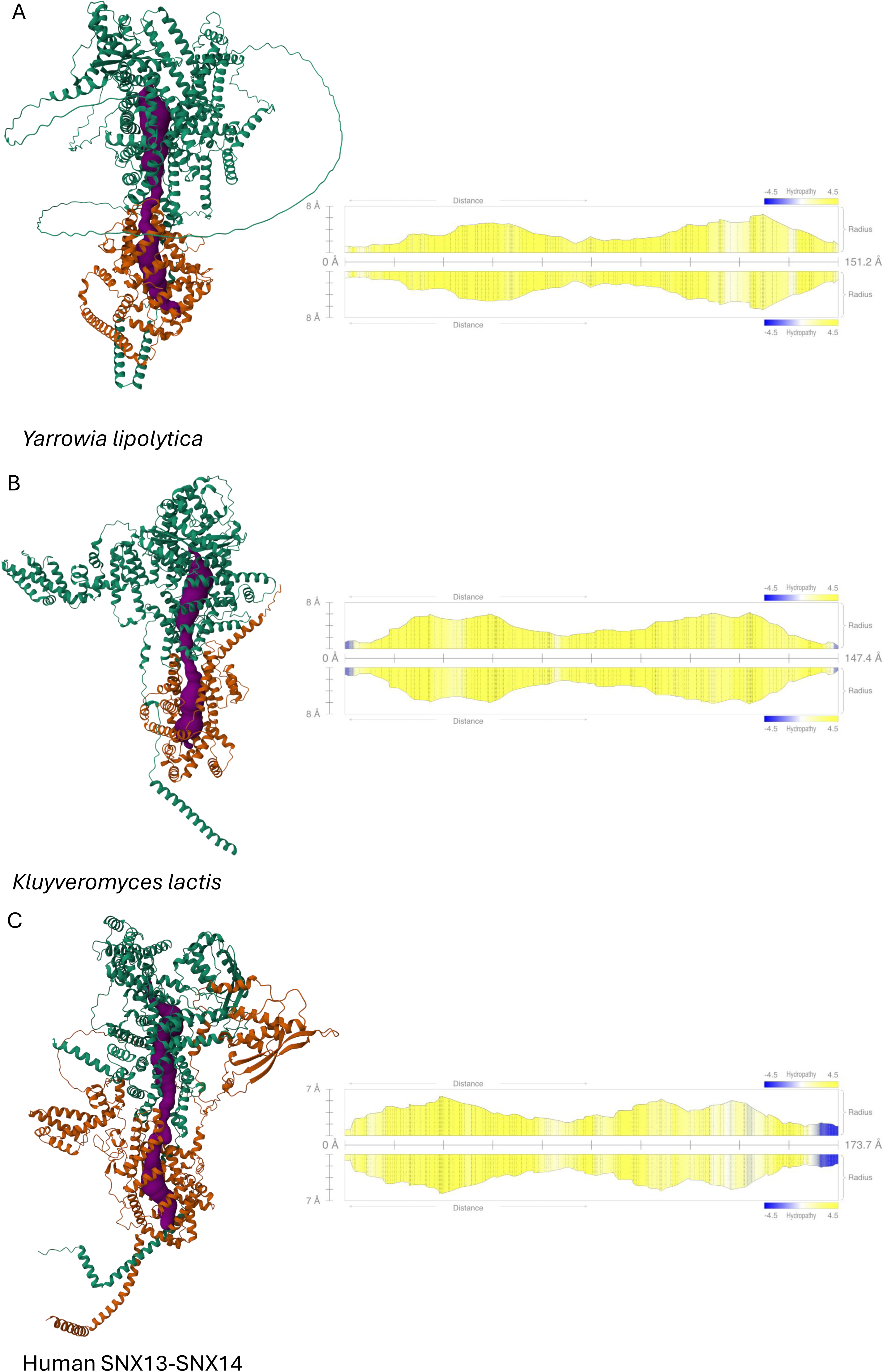
Cross-species structural convergence of a hydrophobic tunnel at the Nvj3-Mdm1 interface. (A-C) AlphaFold-Multimer models of Nvj3-Mdm1 orthologs from *Yarrowia lipolytica* (A) and *Kluyveromyces lactis* (B), and the human paralogous pair SNX13-SNX14 (C). Mdm1 is shown in green and Nvj3 is shown in orange. Predicted tunnel-lining regions identified by MOLEonline are highlighted in purple. Right panels show corresponding tunnel profiles, including radius and hydropathy along the tunnel axis. The interior is predominantly hydrophobic (yellow) and exhibits elongated cavities of lengths ranging between ∼147-173 Å.

Finally, to determine whether this feature extends beyond Saccharomycetes, we examined the human SNX13 and SNX14 as representative metazoan SNX-RGS paralogs related to the Mdm1/Nvj3 family (Henne et al., 2015). Predicted structures of both proteins similarly revealed an elongated internal cavity (173.7 Å) with predominantly hydrophobic character (Figure 6C). Despite increased structural complexity and additional domains, the overall organization of the central channel is maintained. These results suggest that this architecture is likely preserved across species and is consistent with a shared functional role.

To further evaluate whether the predicted human SNX13-SNX14 assembly reflects a broader pairing preference within the metazoan SNX-RGS family, we modeled all possible pairwise combinations among human SNX13, SNX14, SNX19, and SNX25, including four homodimers and six heterodimers. We also modeled all three possible yeast pairings: Nvj3-Mdm1, Mdm1-Mdm1, and Nvj3-Nvj3. These results are summarized in Supplementary Table S4. In yeast, the Nvj3-Mdm1 heterodimer showed higher predicted interface confidence than either homodimer, with a higher ipTM score and lower mean inter-chain PAE. Similarly, among the human SNX-RGS pairings, the SNX13-SNX14 heterodimer showed the strongest AlphaFold interface confidence, whereas most homodimeric models showed lower ipTM scores and/or higher inter-chain PAE values. Although these predictions do not establish biological preference or in vivo complex formation, the comparative pattern indicates that heteromeric assemblies are more strongly supported by AlphaFold confidence metrics than most homomeric alternatives.

## Discussion

The NVJ is a central site for coordinating lipid metabolic pathways, yet the structural organization of its core components has remained poorly defined. Here, we define the Nvj3-Mdm1 complex as a heteromeric assembly that forms a stable interaction interface with features consistent with lipid compatible architecture. Our structural modeling identifies an extended internal conduit spanning ∼175 Å across the Nvj3-Mdm1 interface, with a predominantly hydrophobic lining (Figure 2A). This architecture, not previously described for NVJ tethers, is comparable in diameter and hydropathy to bridge-like lipid transfer proteins (BLTPs) such as Atg2 (Table 2). The convergence of structural, physicochemical, and evolutionary analyses supports a stable assembly with features consistent with a lipid-compatible conduit. Together, these findings support a testable model in which Nvj3-Mdm1 complex may contribute not only to tethering but also to lipid exchange across the NVJ, extending the functional repertoire of membrane contact site tethers beyond simply maintaining membrane proximity (Prinz et al., 2020; Scorrano et al., 2019b). Given the established roles of the NVJ in coordinating lipid metabolism including the regulation of triacylglycerol synthesis and LD storage (Adebayo, Obaseki, Miller, et al., 2026; Hariri et al., 2018, 2019; Kohler & Büttner, 2021), the presence of such a conduit suggests a potential mechanism for spatially organizing lipid flux at this contact site.

The structural organization of the Nvj3-Mdm1 complex suggests a modular architecture that integrates stability with flexibility. A well-defined heterodimeric core, supported by low RMSD values and a sequestered hydrophobic interior, likely serves as a high-affinity interaction scaffold. In contrast, terminal and linker regions exhibit greater conformational variability across the predicted models, suggesting that regions outside the stable interaction core may provide structural flexibility. Such flexibility could be relevant to the spatial organization of the NVJ, although whether it contributes to dynamic membrane spacing remains to be tested experimentally.

Within this framework, the presence of a continuous internal conduit becomes particularly intriguing. Its dimensions and physicochemical properties resemble those of known lipid transfer systems, yet it arises through a fundamentally different structural strategy. Canonical bridge-like lipid transfer proteins (BLTPs) typically form extended hydrophobic grooves within a single polypeptide, often built from β-sheet-rich domains (Maeda et al., n.d.; Osawa et al., 2019; Reinisch & Prinz, 2021). In contrast, the conduit described here emerges from the interface of two interacting proteins, suggesting that similar functional outcomes, namely, the creation of a lipid-compatible pathway, can be achieved through distinct structural solutions. Although direct lipid transfer cannot be inferred from structure alone, the positioning of the tunnel near membrane-proximal regions and its predominantly nonpolar interior are consistent with a lipid-compatible environment.

This concept is further supported by comparison with other contact site systems. In the ERMES complex, lipid transfer is mediated by multiple interacting proteins, including Mmm1 and Mdm12, whose association forms an extended ∼210 Å hydrophobic tunnel capable of accommodating phospholipids (Jeong et al., 2017). Although mechanistically distinct from SMP-domain systems, this example demonstrates that lipid-conducting structures can emerge from protein assemblies rather than single polypeptides. The Nvj3-Mdm1 architecture may represent a related strategy, reinforcing the idea that membrane contact sites employ diverse structural solutions to support lipid exchange.

An additional distinguishing feature of the Nvj3-Mdm1 assembly is that the putative conduit is formed through a heteromeric interface rather than within a single protein. This raises the possibility that lipid handling at the NVJ may be distributed across multiple components, potentially allowing modular regulation of activity, localization, or substrate specificity. In this context, Mdm1 may function as a structural scaffold anchored at the ER, while Nvj3 may contribute to recruitment or regulatory functions. Such division of labor could provide a mechanism for integrating lipid transfer with local metabolic signaling, although further studies will be required to define the specific roles of each component. Consistent with this interpretation, pairwise AlphaFold modeling of all yeast and human SNX-RGS combinations showed that the strongest predicted assemblies were heteromeric rather than homomeric. In yeast, Nvj3-Mdm1 showed substantially higher interface confidence than either Mdm1-Mdm1 or Nvj3-Nvj3. In the human SNX-RGS family, SNX13-SNX14 showed the strongest pairwise confidence among the tested combinations (Supplementary Table S4). These results should be interpreted cautiously, as AlphaFold confidence metrics do not directly measure binding affinity, biochemical favorability, or cellular assembly. Nevertheless, the pattern across all tested pairings supports the idea that heteromeric SNX-RGS assemblies may represent a structurally preferred configuration within this family, warranting future biochemical and cellular validation.

The organization of this system may differ in metazoan lineages that encode fewer SNX-RGS paralogs. For example, *Drosophila* encodes the SNX-RGS protein Snazarus/Snz, which has been shown to associate with peripheral lipid droplets and regulate lipid droplet homeostasis at ER-plasma membrane contact sites in adipocytes (Hariri & Henne, 2022; Ugrankar et al., 2019). This raises an important evolutionary possibility: heteromeric assembly may be favored in lineages with multiple SNX-RGS paralogs, whereas single-paralog systems may maintain lipid-compatible architecture by a single multifunctional protein, homomeric association, or interactions with non-SNX partners. Future comparative modeling and functional analysis across metazoan lineages will be needed to determine whether SNX-RGS family expansion correlates with heteromeric assembly, homomeric association, or single-protein lipid-transfer architectures.

The functional relevance of Nvj3-Mdm1 interface is further supported by its evolutionary conservation. Residues lining both the interaction interface are enriched with highly conserved residues, indicating selective pressure to maintain these structural features (Figure 4). In addition, the observed phylogenetic congruence between Nvj3 and Mdm1 provides evolutionary context for their conserved association across Saccharomycetes (Figure 5). Because this signal likely reflects, at least in part, shared species history rather than direct protein-specific coevolution alone, we interpret the cophylogenetic result cautiously. Together with the enrichment of conserved residues at the predicted interaction interface, these findings support the view that the Nvj3-Mdm1 association has been maintained within a conserved functional framework, rather than representing a purely incidental structural interaction.

Several important questions remain to be addressed. The present study relies on computational structural predictions, and experimental validation will be required to determine whether the predicted conduit supports lipid transport *in vivo*. The conformational flexibility suggested by PAE and RMSD analyses (Figure 1B, C) indicates that the relative orientation of Nvj3 and Mam1 may vary, which could influence tunnel accessibility and function. Future biochemical and reconstitution assays will be necessary to assess lipid specificity, directionality, and transport capacity. In addition, coarse-grained molecular dynamics simulations could provide insight into lipid accessibility and the dynamic behavior of the conduit. More broadly, understanding how this interface contributes to NVJ function may reveal new principles of membrane contact site biology. Given the central role of these structures in fungal physiology, it will also be of interest to explore whether this assembly represents a tractable target for modulating lipid-associated processes.

## Methods

### Structural Analysis and Visualization

Structural prediction of the Nvj3-Mdm1 complex by AlphaFold3 was performed using the AlphaFold Server (https://alphafoldserver.com/), using amino acid sequences of Mdm1 (YML104C) and Nvj3 (YDR179W-A) retrieved from the Saccharomyces Genome Database (SGD; <u>(Reinisch and Prinz, 2021; Osawa et al., 2019; Maeda et al.)</u>). Five ranked models (Models 0-4) were analyzed within the UCSF ChimeraX v1.6 (Pettersen et al., 2021). The model with the highest ranking was used for all downstream analysis. Structural superimposition was performed using the Matchmaker (mmaker) command-line tool, utilizing the Needleman-Wunsch algorithm for pairwise sequence alignment with the BLOSUM-62 residue similarity matrix and secondary structure weighting (SS fraction = 0.3; gap penalties: HH/SS/other = 18/18/6; gap extension = 1). Structural divergence was quantified by calculating the per-residue Root Mean Square Deviation (RMSD) relative to Model 0, generating a seq_rmsd attribute for the entire ensemble. The biochemical environment of this stabilized core was further characterized by generating molecular surfaces mapped for hydrophobicity using the *mlp* command and Coulombic electrostatic potential was calculated using the *coulombic* command in ChimeraX. A side-view cross-section of the interface was created using the clipping tool to expose the interior of the central binding cavity.

Domain boundaries used for structural coloring were identified using HHpred through the Bioinformatics Toolkit server.

### Binding affinity and interface analysis using PRODIGY

Binding affinity of Mdm1 and Nvj3 interaction was estimated using PRODIGY https://bianca.science.uu.nl/prodigy/ (Xue et al., 2016). The AlphaFold-Multimer model was used as input with chains corresponding to Mdm1 and Nvj3 defined as interacting partners. Calculations were performed at 25 °C, and intermolecular contact analysis was enabled to quantify contact types and interface composition, including apolar-apolar and charged-apolar interactions.

### Tunnel Detection and Visualization

Tunnel detection within the Nvj3-Mdm1 complex was performed using MOLEonline 2.5 server (https://mole.upol.cz/) (Pravda et al., 2018) in “channels” mode with optimized parameters to capture the internal conduit (probe radius 9.17 Å; interior threshold 0.87). The VoronoiScale weight function was applied to map the tunnel geometry, maintaining a bottleneck tolerance of 3. To ensure the chemical environment of the tunnel was accurately represented, hydrogens were included in the calculation. The resulting json file informed the subsequent visualization in ChimeraX v1.9 (Pettersen et al., 2021). The structure was oriented to align the tunnel approximately with the Z-axis, and clipping planes were applied via the Side View tool to expose the longitudinal channel through the complex. Surface capping was enabled to clearly visualize the interior morphology, and hydrophobicity coloring was applied to highlight the chemical nature of the tunnel walls.

### Comparative Tunnel Analysis

Comparative analyses were conducted using predicted structures of Atg2, Fmp27, and Hob2 from *Saccharomyces cerevisiae* retrieved from the AlphaFold Protein Structure Database (https://alphafold.ebi.ac.uk/) Atg2: AF-P53855-F1; Fmp27: AF-Q06179-F1; Hob2: AF-Q06116-F1). Tunnel detection and quantitative characterization for all analyzable structures were performed using MOLEonline 2.5 server (https://mole.upol.cz/) with identical parameters used for Nvj3-Mdm1. For each detected tunnel, MOLEonline calculates geometric properties at discrete segments along the tunnel centerline. Average radius values were derived from the probe-defined (b-radius) profile, where the b-radius represents the radius of the largest sphere that can pass through the channel without steric clashes, and the corresponding diameter was estimated as twice this value.

### Evolutionary Conservation Analysis

Residue-level evolutionary conservation was calculated using ConSurf (Ashkenazy et al., 2016) via a Google Colab implementation. No custom multiple sequence alignment was provided; therefore, ConSurf automatically identified homologous sequences from UniRef100 and generated a protein-specific MSA for each chain. ConSurf estimates the evolutionary conservation of each amino acid position by aligning homologous sequences, reconstructing their phylogenetic relationships, and calculating position-specific evolutionary rates, where slowly evolving residues are assigned higher conservation scores. Conservation scores were mapped onto the corresponding structural models, and the individual chains were subsequently combined and visualized in ChimeraX.

ConSurf output files for Mdm1 and Nvj3 were parsed in R (RStudio Team 2024) using a custom pipeline to extract residue-level conservation scores and structural mapping information. To ensure robustness, residues were filtered to retain only positions with valid structural mapping, no low-confidence annotation, and sufficient representation in the multiple sequence alignment (≥ 6 sequences).

Interface residues were defined from PRODIGY-derived residue-residue contact files by extracting unique interacting positions from Mdm1 and Nvj3. Visualization in ChimeraX was used to refine this set by excluding spatially peripheral contacts, retaining only residues contributing to the central interaction interface. To quantify conservation at the interface, residues were classified as interface or non-interface, and the proportion of highly conserved residues with grades 8-9 in each category was calculated across the combined dataset and separately for each protein. Statistical enrichment of conserved residues at the interface was assessed using Fisher’s exact test.

### Sequence Collection and Alignment

Protein sequences for Mdm1 (Group 96377at4891) and Nvj3 (Group 113872at4891) were collected at the Saccharomycetes taxonomic level from OrthoDB v12.1database (Kuznetsov et al., 2022). Raw FASTA files were parsed in RStudio (RStudio Team 2024) extracting gene identifiers and organism names from headers. Species names were standardized to genus and species, and for species with multiple sequence entries, only the longest sequence was retained. From these curated datasets, 120 species were shared between Mdm1 and Nvj3, and only sequences from these species were retained for downstream analyses. Curated sequences were exported in FASTA format and aligned using MAFFT v7 with default parameters (Katoh et al., 2019) to generate multiple sequence alignments for each protein.

### Cophylogenetic Analysis

Phylogenetic trees for Mdm1 and Nvj3 were separately reconstructed using Bayesian inference in MrBayes v3.2 under a general time-reversible substitution copjmodel with a proportion of invariable sites and gamma-distributed rate variation across sites Markov chain Monte Carlo (MCMC) analyses were run for 5,000,000 generations with sampling every 100 generations using two chains. Convergence was assessed using standard diagnostics, and a majority-rule consensus tree with posterior probabilities was generated after discarding burn-in. Cophenetic distance matrices were calculated from the trees, and a one-to-one association matrix representing the shared species was constructed. Co-evolutionary patterns were assessed using PACo (paco R package) with 1,000 permutations, following principal coordinate transformation with Cailliez correction to account for non-Euclidean distances.

## Acknowledgements

We thank Dr. Maya Schuldiner and Ms. Maria-Del-Rosario Valenti for insightful discussions and optimizations of early AF predications. H. Hariri is supported by the National Institutes of Health National Institute of General Medical Sciences (R35GM150892), Wayne State University start-up funds and University Research Award, Karmanos Cancer Institute - Strategic Research Initiative Grant (KCI-SRIG), Richard Barber Interdisciplinary Research Program.

## Author contributions

M. Aboumourad and H. Hariri, conception and design of the study, data interpretation, and revising the manuscript. M. Aboumourad experimental design, data generation and analysis, manuscript drafting and figure making.

## Availability of data and materials

This study used publicly available protein sequences and structural data. Protein sequences were retrieved from OrthoDB, and reference structures or models used for comparative analyses were obtained from the AlphaFold Protein Structure Database, as described in the Methods. The datasets generated during the current study, including AlphaFold Server models, MOLEonline tunnel outputs, conservation analysis results, phylogenetic trees, PACo outputs, processed figure data, and R scripts used for phylogenetic, cophylogenetic, and statistical analyses, are available upon request.

## Supplementary Material

**Supplementary Figure 1. Domain mapping of the Nvj3-Mdm1 heterodimer.** Predicted Nvj3-Mdm1 heterodimer shown in cartoon representation with selected regions colored and the remaining structure shown in grey. In Mdm1, the transmembrane domain is colored cyan, PXA orange, PX blue, and PXC green. In Nvj3, PXA is colored purple and PXC is colored red. Domain boundaries predicted by HHpred are: Mdm1: transmembrane helix, residues 1-45; PXA, residues 87-271; PX, residues 763-900; and PXC, residues 993-1106. Nvj3: PXA, residues 98-238; and PXC, residues 348-449.

**Supplementary Figure 2. AlphaFold3 prediction of the Nvj3-Mdm1 complex in the presence of oleic acid for visualization reveals OLA positioned within the internal cavity.**

**A)** Overview of the predicted Nvj3-Mdm1 heterodimer generated using AlphaFold3 in the presence of 50 copies of oleic acid (OLA). Nvj3 is colored pink, Mdm1 is colored steel blue, and OLA molecules are shown as sticks within the elongated internal cavity.

**B)** Zoomed-in view of the proximal, nuclear ER-facing entrance of the OLA-occupied cavity.

**C)** View rotated 180° relative to panel B, showing the opposite distal side of the cavity, which appears more constricted than the proximal entrance.

**Table S1. Structural similarity among AlphaFold-predicted Nvj3-Mdm1 models.**

Pairwise structural comparisons were performed in ChimeraX MatchMaker by aligning four AlphaFold-predicted Nvj3-Mdm1 models to the top-ranked reference model. RMSD values are reported separately for each chain, with chain A corresponding to Mdm1 and chain B corresponding to Nvj3. Pruned RMSD values reflect the aligned structural core after excluding poorly matching regions, whereas total RMSD values include all aligned residue pairs. Low pruned RMSD values indicate that the predicted core folds are consistent across models, while higher total RMSD values likely reflect variation in flexible or low-confidence regions.

**Table S2. Residues lining the predicted Nvj3-Mdm1 tunnel.**

Residues lining the internal tunnel of the predicted Nvj3-Mdm1 complex as identified in the MOLEonline tunnel output. The table lists each tunnel-lining residue by amino acid, sequence position, and protein chain, with chain A corresponding to Mdm1 and chain B corresponding to Nvj3.

**Table S3. Interface residues identified in the predicted Nvj3-Mdm1 complex.**

Interface residues were defined from the predicted Nvj3-Mdm1 structural model and are listed by protein, residue identity and position, and ConSurf conservation score. The complete interface residue set is provided to support the selectively labeled inset in Figure 4, where only residues with a score of 9 are labeled for readability.

**Table S4. Pairwise AlphaFold modeling of yeast and human SNX-RGS combinations.**

Pairwise AlphaFold3 modeling was performed for all possible yeast and human SNX-RGS combinations. Yeast comparisons included Mdm1-Nvj3, Mdm1-Mdm1, and Nvj3-Nvj3. Human comparisons included all four homodimers and six heterodimers among SNX13, SNX14, SNX19, and SNX25. For each pair, pTM, ipTM, mean inter-chain PAE, and presence of an elongated predicted structure are reported. Higher ipTM and lower mean inter-chain PAE indicate greater predicted interface confidence. Nvj3-Mdm1 and SNX13-SNX14 showed the strongest predicted interface confidence among the yeast and human pairings, respectively, with the highest ipTM scores and lowest mean inter-chain PAE values.

