## Supplementary figures and images for "Structural Organization of the Nvj3-Mdm1 Complex Reveals a Conserved Lipid-Compatible Contact Site Module"

### Supplemental Figures

Supplementary Figure 1

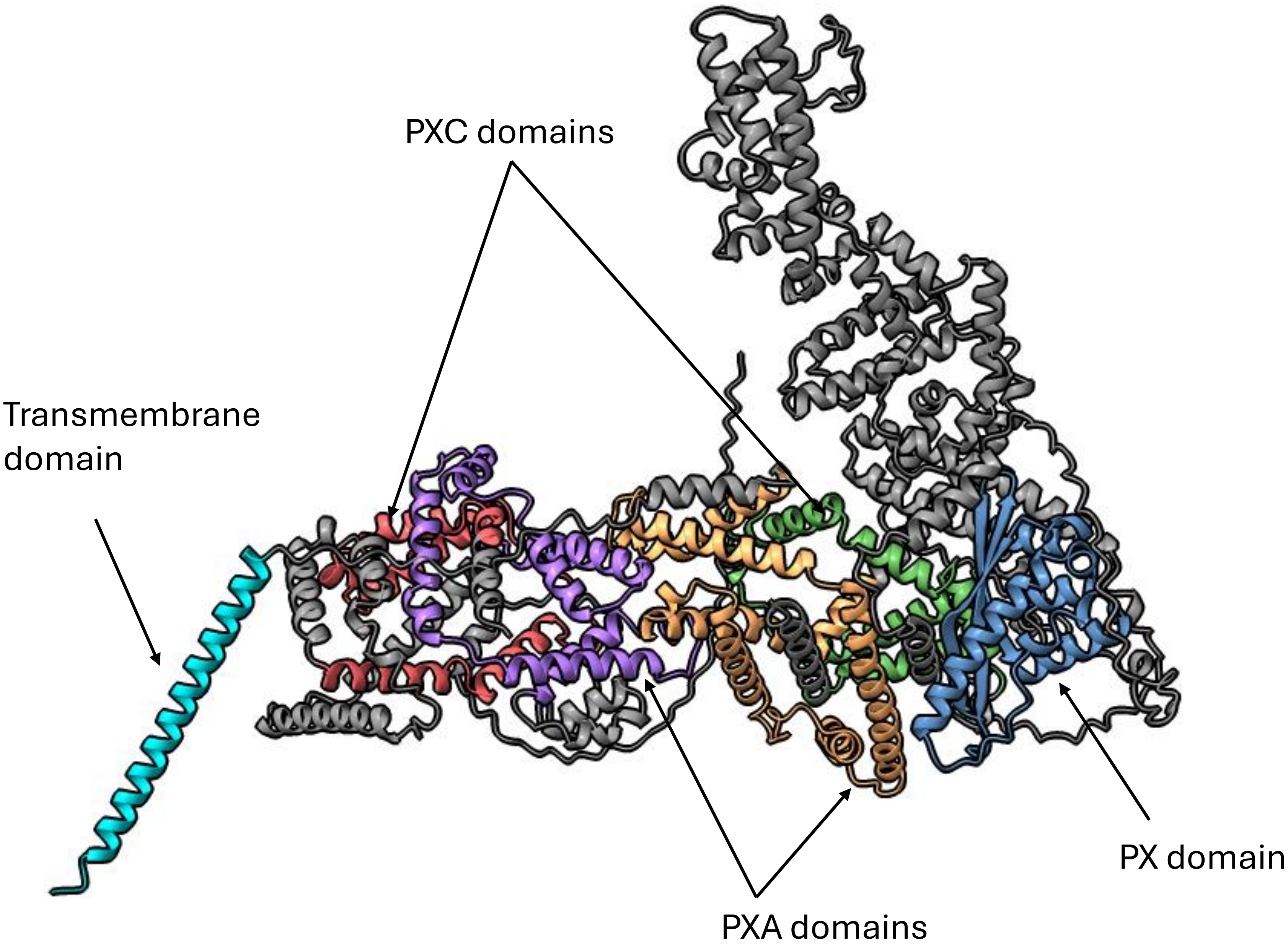

Supplementary Figure 2

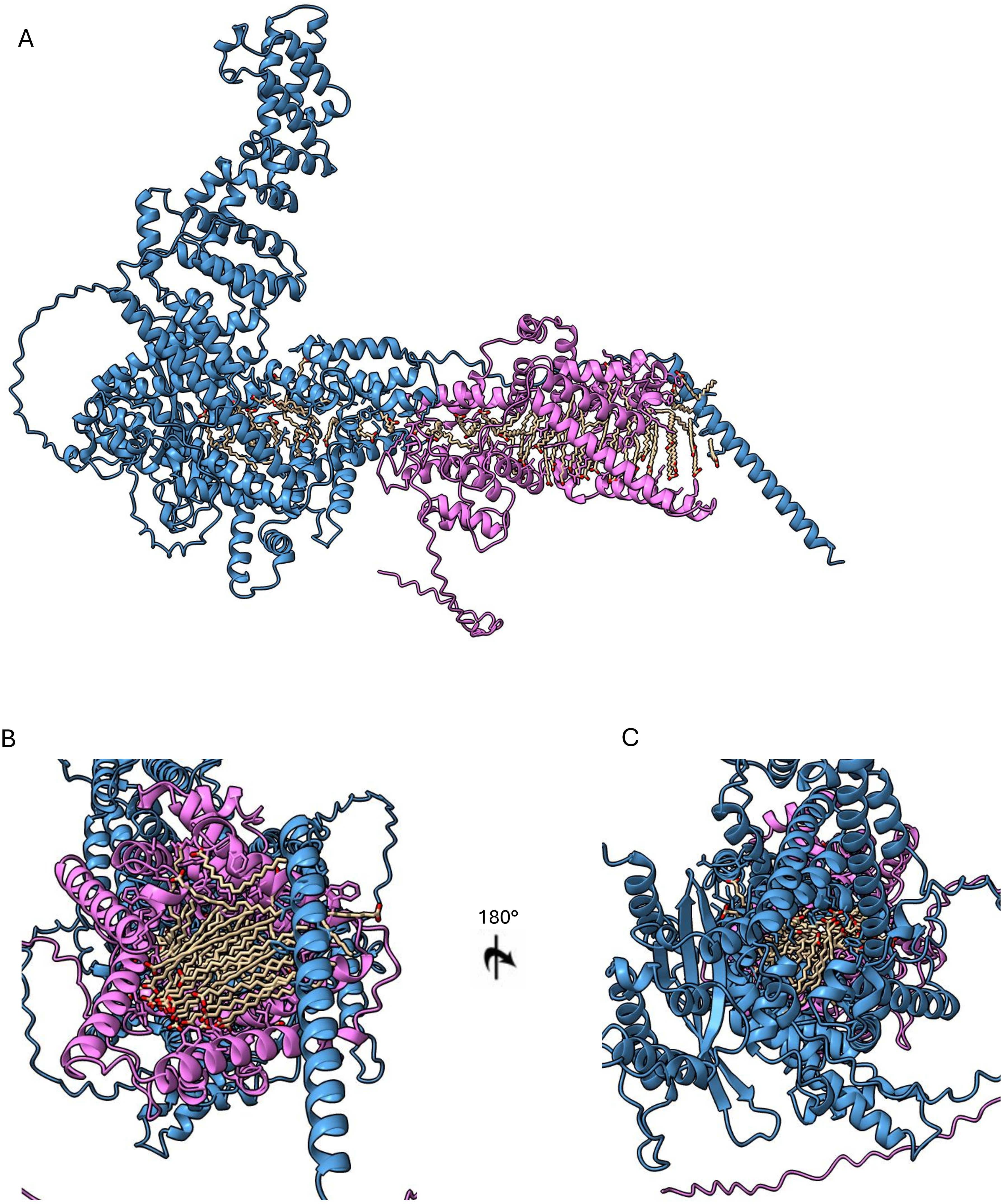
