## Supplemental Tables for "Structural Organization of the Nvj3-Mdm1 Complex Reveals a Conserved Lipid-Compatible Contact Site Module"

**Table S1. Structural similarity among AlphaFold-predicted Nvj3-Mdm1 models.** Pairwise structural comparisons were performed in ChimeraX MatchMaker by aligning four AlphaFold-predicted Nvj3-Mdm1 models to the top-ranked reference model. RMSD values are reported separately for each chain, with chain A corresponding to Mdm1 and chain B corresponding to Nvj3. Pruned RMSD values reflect the aligned structural core after excluding poorly matching regions, whereas total RMSD values include all aligned residue pairs. Low pruned RMSD values indicate that the predicted core folds are consistent across models, while higher total RMSD values likely reflect variation in flexible or low-confidence regions.

| Match Model | Chain | Pruned RMSD (Å) | Pruned Pairs | Total RMSD (Å) | Total Pairs | Score |
|---|---|---|---|---|---|---|
| **Model 1** | A | 1.091 | 876 | 5.612 | 1127 | 5672.7 |
| **Model 2** | A | 0.690 | 693 | 8.027 | 1127 | 5636.1 |
| **Model 3** | A | 0.784 | 817 | 8.728 | 1127 | 5703.3 |
| **Model 4** | A | 0.686 | 672 | 12.562 | 1127 | 5676.9 |
| **Model 1** | B | 0.523 | 389 | 10.299 | 463 | 2390.6 |
| **Model 2** | B | 0.505 | 386 | 22.129 | 463 | 2369.0 |
| **Model 3** | B | 0.438 | 400 | 11.158 | 463 | 2352.2 |
| **Model 4** | B | 0.515 | 386 | 9.251 | 463 | 2385.2 |

**Table S2. Residues lining the predicted Nvj3-Mdm1 tunnel.** Residues lining the internal tunnel of the predicted Nvj3-Mdm1 complex as identified in the MOLEonline tunnel output. The table lists each tunnel-lining residue by amino acid, sequence position, and protein chain, with chain A corresponding to Mdm1 and chain B corresponding to Nvj3.

| Residue | Position | Chain |
|---|---|---|
| ILE | 99 | A |
| VAL | 104 | A |
| TRP | 107 | A |
| PHE | 119 | A |
| ILE | 123 | A |
| LEU | 127 | A |
| LEU | 131 | A |

| | | |
|---|---|---|
| VAL | 134 | A |
| LEU | 138 | A |
| ILE | 147 | A |
| LEU | 151 | A |
| LEU | 237 | A |
| VAL | 238 | A |
| LEU | 241 | A |
| MET | 242 | A |
| VAL | 245 | A |
| LEU | 246 | A |
| ILE | 250 | A |
| LEU | 254 | A |
| TRP | 264 | A |
| ARG | 267 | A |
| LEU | 271 | A |
| TYR | 275 | A |
| LYS | 279 | A |
| PHE | 649 | A |
| LEU | 661 | A |
| PHE | 662 | A |
| LYS | 750 | A |
| ARG | 753 | A |
| THR | 754 | A |
| LYS | 757 | A |
| VAL | 1002 | A |
| GLN | 1005 | A |
| LEU | 1006 | A |
| TYR | 1014 | A |
| LEU | 1022 | A |
| ALA | 1031 | A |
| PHE | 1035 | A |
| ARG | 1084 | A |
| VAL | 1085 | A |
| ILE | 1119 | A |
| PHE | 104 | B |
| ILE | 105 | B |
| ILE | 108 | B |
| PHE | 109 | B |
| PHE | 128 | B |

| | | |
|---|---|---|
| LEU | 132 | B |
| LEU | 135 | B |
| VAL | 136 | B |
| LEU | 139 | B |
| LEU | 143 | B |
| LEU | 154 | B |
| ILE | 158 | B |
| LEU | 162 | B |
| LEU | 212 | B |
| GLN | 213 | B |
| PHE | 216 | B |
| LEU | 217 | B |
| PHE | 220 | B |
| VAL | 225 | B |
| PHE | 230 | B |
| ILE | 233 | B |
| LEU | 239 | B |
| ILE | 246 | B |
| ILE | 250 | B |
| LYS | 281 | B |
| MET | 285 | B |
| LEU | 288 | B |
| PHE | 316 | B |
| PHE | 319 | B |
| LEU | 333 | B |
| ILE | 336 | B |
| CYS | 337 | B |
| LEU | 340 | B |
| ILE | 361 | B |
| LYS | 365 | B |
| ILE | 366 | B |
| LEU | 375 | B |
| PHE | 376 | B |
| LEU | 379 | B |
| TYR | 420 | B |
| LEU | 422 | B |
| ILE | 425 | B |
| ILE | 451 | B |
| ILE | 454 | B |

| | | |
|---|---|---|
| ILE | 455 | B |
| LEU | 661 | A |
| VAL | 1002 | A |
| GLN | 1005 | A |
| LEU | 1006 | A |
| ASP | 1080 | A |
| THR | 1081 | A |
| GLN | 213 | B |
| ILE | 336 | B |
| CYS | 337 | B |

**Table S3. Interface residues identified in the predicted Nvj3-Mdm1 complex.**

Interface residues were defined from the predicted Nvj3-Mdm1 structural model and are listed by protein, residue identity and position, and ConSurf conservation score. The complete interface residue set is provided to support the selectively labeled inset in Figure 4, where only residues with a score of 9 are labeled for readability.

| Protein | Residue ID | Conservation Score |
|---|---|---|
| Mdm1 | W107 | 8 |
| Mdm1 | F108 | 8 |
| Mdm1 | K110 | 5 |
| Mdm1 | I111 | 8 |
| Mdm1 | D112 | 8 |
| Mdm1 | A117 | 7 |
| Mdm1 | E118 | 1 |
| Mdm1 | F119 | 9 |
| Mdm1 | V122 | 4 |
| Mdm1 | I123 | 7 |
| Mdm1 | W125 | 1 |
| Mdm1 | R126 | 3 |
| Mdm1 | K210 | 3 |
| Mdm1 | E213 | 6 |
| Mdm1 | R217 | 4 |
| Mdm1 | I221 | 5 |
| Mdm1 | D231 | 4 |
| Mdm1 | E232 | 9 |
| Mdm1 | L233 | 9 |
| Mdm1 | D234 | 5 |

| | | |
|---|---|---|
| Mdm1 | S235 | 9 |
| Mdm1 | L236 | 5 |
| Mdm1 | L237 | 5 |
| Mdm1 | V238 | 9 |
| Mdm1 | T240 | 6 |
| Mdm1 | L241 | 8 |
| Mdm1 | M242 | 7 |
| Mdm1 | E244 | 9 |
| Mdm1 | V245 | 8 |
| Mdm1 | T248 | 6 |
| Mdm1 | C249 | 6 |
| Nvj3 | K127 | 4 |
| Nvj3 | F128 | 8 |
| Nvj3 | E131 | 9 |
| Nvj3 | L132 | 5 |
| Nvj3 | L135 | 5 |
| Nvj3 | K192 | 7 |
| Nvj3 | Y194 | 8 |
| Nvj3 | M196 | 2 |
| Nvj3 | L197 | 3 |
| Nvj3 | E200 | 3 |
| Nvj3 | M206 | 3 |
| Nvj3 | S207 | 6 |
| Nvj3 | K209 | 4 |
| Nvj3 | S210 | 9 |
| Nvj3 | L211 | 5 |
| Nvj3 | L212 | 8 |
| Nvj3 | Q213 | 9 |
| Nvj3 | S215 | 8 |
| Nvj3 | F216 | 6 |
| Nvj3 | D218 | 9 |
| Nvj3 | S219 | 8 |
| Nvj3 | E223 | 4 |
| Nvj3 | L224 | 8 |

**Table S4. Pairwise AlphaFold modeling of yeast and human SNX-RGS combinations.**

Pairwise AlphaFold3 modeling was performed for all possible yeast and human SNX-RGS combinations. Yeast comparisons included Mdm1-Nvj3, Mdm1-Mdm1, and Nvj3-

Nvj3. Human comparisons included all four homodimers and six heterodimers among SNX13, SNX14, SNX19, and SNX25. For each pair, pTM, ipTM, mean inter-chain PAE, and presence of an elongated predicted structure are reported. Higher ipTM and lower mean inter-chain PAE indicate greater predicted interface confidence. Nvj3-Mdm1 and SNX13-SNX14 showed the strongest predicted interface confidence among the yeast and human pairings, respectively, with the highest ipTM scores and lowest mean inter-chain PAE values.

| System | Protein pair | Pair type | pTM | ipTM | Mean inter-chain PAE (Å) | Elongated Structure |
|---|---|---|---|---|---|---|
| Yeast | Mdm1-Nvj3 | Heterodimer | 0.64 | 0.64 | 5.06 | Yes |
| Yeast | Mdm1-Mdm1 | Homodimer | 0.36 | 0.21 | 18.44 | Yes |
| Yeast | Nvj3-Nvj3 | Homodimer | 0.45 | 0.33 | 13.66 | No |
| Human | SNX13-SNX13 | Homodimer | 0.54 | 0.47 | 8.65 | Yes |
| Human | SNX14-SNX14 | Homodimer | 0.37 | 0.25 | 21.57 | No |
| Human | SNX19-SNX19 | Homodimer | 0.4 | 0.32 | 18.82 | Yes |
| Human | SNX25-SNX25 | Homodimer | 0.41 | 0.34 | 11.59 | Yes |
| Human | SNX13-SNX14 | Heterodimer | 0.64 | 0.64 | 6.52 | Yes |
| Human | SNX13-SNX19 | Heterodimer | 0.53 | 0.51 | 18.82 | Yes |
| Human | SNX13-SNX25 | Heterodimer | 0.49 | 0.44 | 11.08 | Yes |
| Human | SNX14-SNX19 | Heterodimer | 0.48 | 0.47 | 17.49 | Yes |
| Human | SNX14-SNX25 | Heterodimer | 0.47 | 0.47 | 19.31 | Yes |
| Human | SNX19-SNX25 | Heterodimer | 0.45 | 0.44 | 14.80 | Yes |
